# PINK1 knockout rats show premotor cognitive deficits measured through a complex maze

**DOI:** 10.1101/2024.01.18.576285

**Authors:** Isabel Soto, Vicki A. Nejtek, David P. Siderovski, Michael F. Salvatore

## Abstract

Cognitive decline in Parkinson’s disease (PD) emerges up to 10 years before clinical recognition. Neurobiological mechanisms underlying premotor cognitive impairment in PD can potentially be examined in the PINK1^-/-^ rat, which exhibits a protracted motor onset. To enhance translation to human PD cognitive assessments, we tested a modified multiple T-maze, which measures cognitive flexibility similarly to the Trail-Making Test in humans. Like human PD outcomes, PINK1^-/-^ rats made more errors and took longer to complete the maze than wild types. Thus, we have identified a potential tool for assessing cross-species translation of cognitive functioning in an established PD animal model.

---

Parkinson’s disease (PD) is primarily known as a motor disease resulting from dopamine (DA) depletion in the nigrostriatal pathway. However, a number of non-motor symptoms (NMS) also significantly impact a PD patient’s quality of life [1]. One such NMS is cognitive impairment, affecting close to one-third of PD patients, and presenting up to a decade prior to the onset of motor decline. These cognitive impairments can manifest as a subtle decline in executive functioning and typically evade detection by global cognitive tests [2-4]. Gaining an understanding of the mechanisms underlying prodromal executive function impairments in PD will help identify strategies to delay the onset of motor impairment, particularly as PD patients typically seek medical attention only well after motor symptoms first appear [5].

Phosphatase and tensin homolog-induced kinase 1 (PINK1) is a protein involved in mitophagy; a loss of function mutation within PINK1 is associated with early-onset PD [6]. The PINK1^-/-^ rat, a genetic preclinical model of early-stage PD, may provide an opportunity to evaluate cognitive impairments that manifest during the premotor phase of PD. Still in its infancy, studies using the PINK1^-/-^ model show some controversy about the age of onset of symptoms and the extent of these deficits across the lifespan of the rat model [7-11]. Apart from motor dysfunction, one lab has reported vocalization deficits in PINK1^-/-^ rats [9], and only one study to-date has found object discrimination deficits at 5-months-old using the novel object recognition (NOR) task as a premotor feature in this model [11]. We reasoned that a more precise characterization of executive functioning in the PINK1^-/-^ rat could present an important experimental procedure toward establishing its suitability for investigating early biological markers and mechanisms related to cognitive impairment in PD.

The Trail-Making Test (TMT) is a goal-directed task used to measure attention, cognitive flexibility, and working memory in humans [4,13]. Our primary goal was to explore whether a rodent task might detect premotor executive functioning deficits that would translate better to TMT outcomes in human PD than other animal testing paradigms [14-15]. To accomplish this goal, we modified the Cincinnati Water Maze (CWM), and investigated its ability to provide translatable outcomes from genetic PD rats that could mirror those from humans with PD. The CWM measures goal-directed activity and spatial navigation processes that rely on visuospatial function, motivation, and episodic memory associated with the striatum, prefrontal cortex, and hippocampus [16-18]. Targeting these neural domains of cognition increases the probability that the preclinical (rodent-based) outcomes would align with established TMT results in humans with PD. In doing so, we expected to measure episodic memory, and visuospatial abilities that are more sensitive to subtle cognitive decline (like the TMT) than those measuring global cognitive functioning [4, 19-23].

Here, we modified the CWM to a non-water appetitive 5-T choice-arm (5-T) maze to avoid the anxiety-inducing stress response in rats triggered by swimming that introduces an inherent confound to cognitive outcomes [16, 24]. Likewise, we chose a modest food restriction of 20% from daily intake during the testing period rather than more severe restrictions for an extended period of time, the latter which could confound interpretations of cognitive performance. Studies of cognition under severe food restriction show mixed results. For example, some studies have shown that even an acute period of food restriction can significantly interfere with cogntivie functioning, but other studies show an improvement [25-26]. Heavily restricting food intake to a degree where body mass is afffected could negatively impact cognition or drive some animals to seek the food reward more than others, thus adversely influencing the results [25]. Therefore, we administered an appetitive 5-T maze to 4-month-old PINK1^-/-^ rats with mitigated confounding variable to better reflect the premotor cognitive decline observed in early-stage PD human subjects [4, 19].

Thirty-four 4-month-old male Long Evans rats (PINK1^-/-^ n=20, Wild-type [WT] n=14) were assessed on latency and number of errors during the 5-T maze. Two rats were unable to find the endpoint after 5 min on day 1, and only 1 rat failed in this fashion on day 4 of testing. PINK1^-/-^ rats took a significantly longer time to complete the maze (Fig. 1A). and committed significantly more errors compared to WT rats (Fig. 1B). We did not identify a significant effect on trial day performance in either the WT or PINK1^-/-^ group (Genotype (F(1,32)= 5.01, ^*^*p*= 0.03), Trial Day (F(4,121)= 1.01, not significant), Genotype x Trial Day (F(4, 121)= 1.19, not significant). Considering PINK1^-/-^ rats eventually develop motor dysfunction [7-9] spontaneous locomotion was evaluated using the open-field test (OFT) to address the possibility that the longer navigation time in the PINK1^-/-^ rats was related to decreased locomotor activity. Rather than a decrease, PINK1^-/-^ rats displayed hyperactivity during both pre- and post-cognitive testing compared to WT rats, with significantly greater distance traveled in the open field (Fig. 2A) and higher average speed (Fig. 2B) compared to WT rats in the first 5 min or a full hour of OFT.

**Fig 1A.**
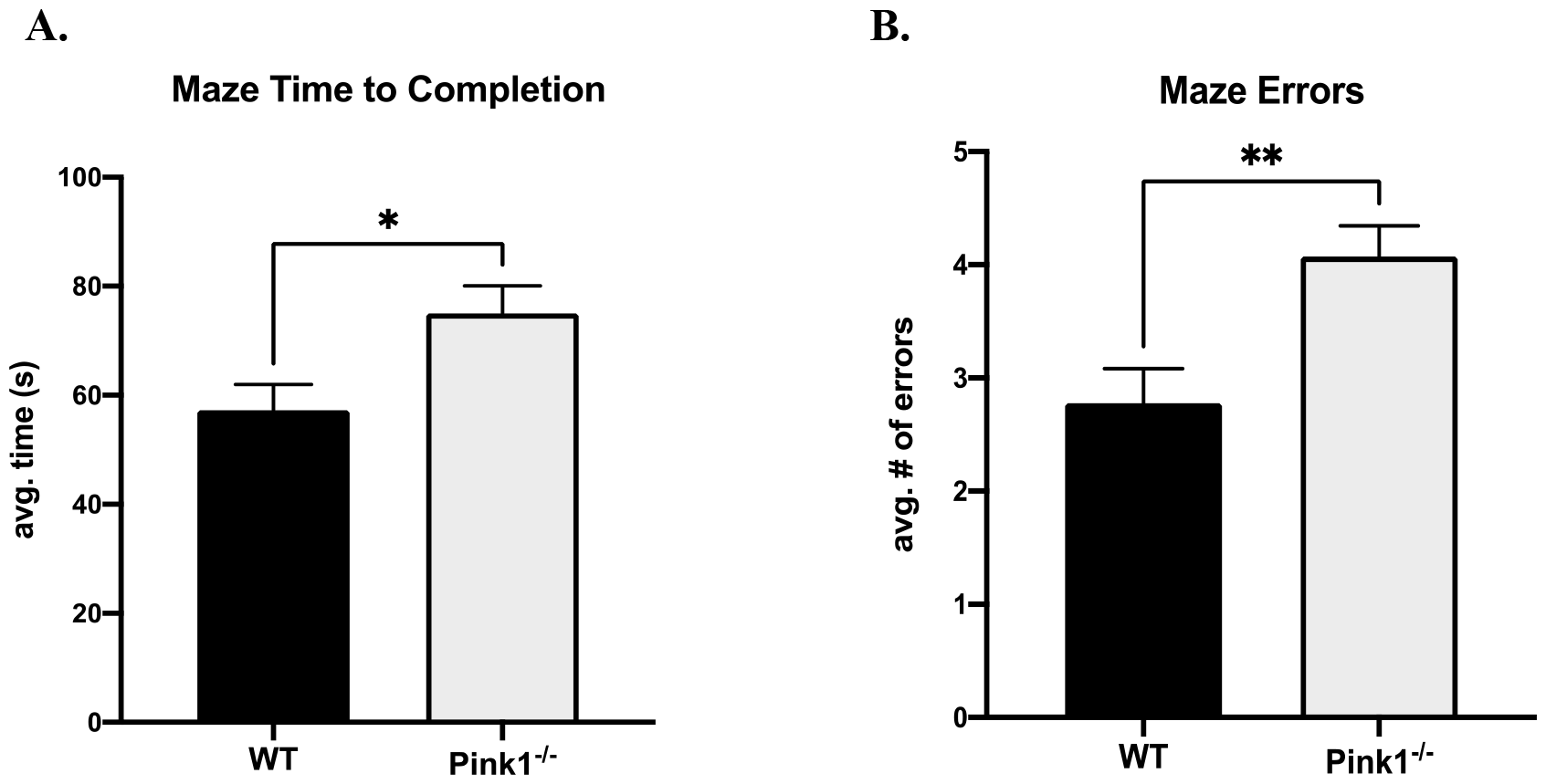
**Mean Maze Time to Completion**. Pink1^-/-^ rats on average took a significantly longer time to complete the maze than WT controls (t=2.10, ^*^*p*=0.04, df=31). **Fig1.B. Mean Maze Errors**. Pink1 KO rats made significantly more errors in completing the maze than WT controls (t=3.13, ^**^*p*=0.003, df=32).

**Fig 2.A.**
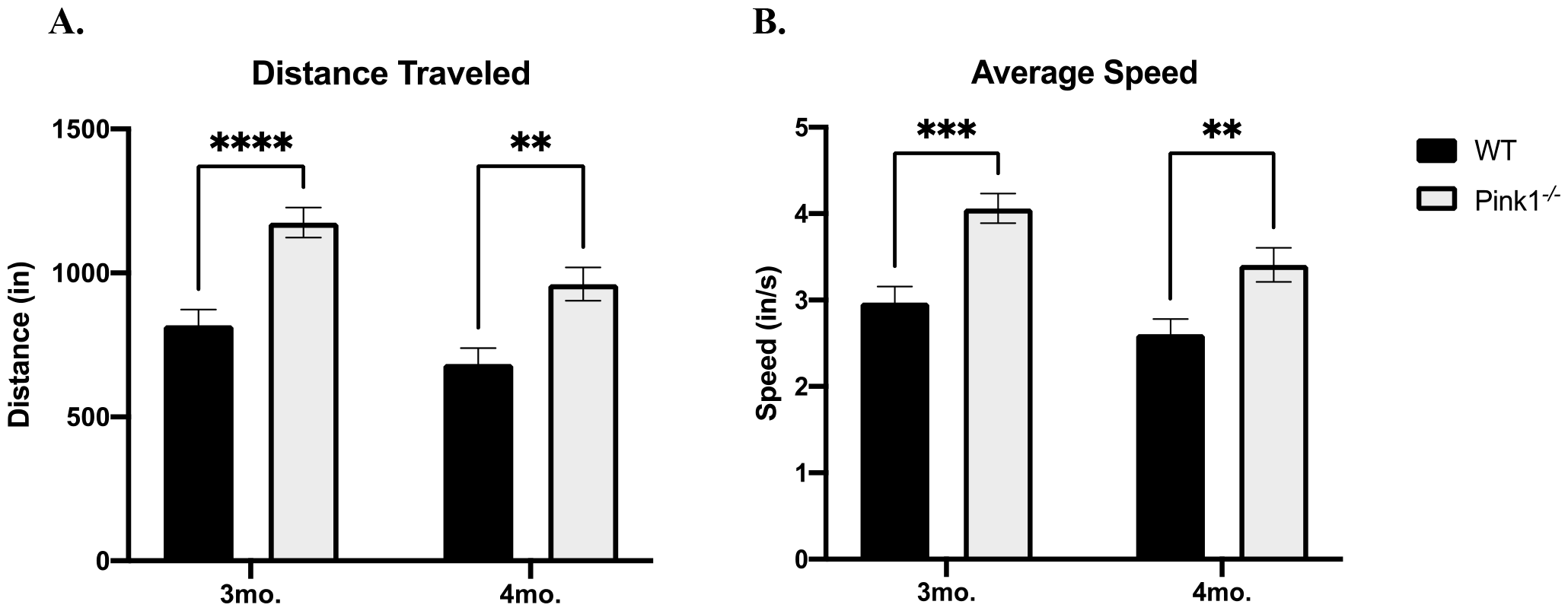
**Distance Traveled in 5 min**. Locomotor activity was significantly greater in Pink1^-/-^ versus WT rats at both 3 and 4mo. Genotype (F(1,31)= 20.67, ^****^*p* < 0.0001), Age (F(1,31)= 20.73, ^****^*p* < 0.0001), Genotype x Age (F(1,31)= 1.09, ns). **Fig2.B. Average Speed in 5 min**. Pink1^-/-^ rats also had a significantly faster average speed versus WT at both 3 and 4mo. Genotype (F(1,31)= 16.18, ^***^*p=*0.0003), Age (F(1,31)= 17.21, ^***^*p=*0.0002). Genotype x Age (F(1,31)= 1.37, ns).

Our results reveal that PINK1^-/-^ rats take a significantly longer time to complete the maze, despite hyperkinetic OFT outcomes. They also make significantly more errors than WT rats. These cognitive outcomes align with prior investigations in PINK1^-/-^ rodents, which exhibit deficits in memory and learning capacities [11, 27]. The 5-T maze requires rats to recall which paths they must travel from the beginning to the end with minimal errors throught the “five choice arms” configuration. Human PD subjects in the TMT test must also use working memory to recall where randomly-placed letters and numbers are located, with the goal to complete the task as quickly as possible. Therefore, this multiple T-maze results has greater translation to human PD assessments than the more commonly used NOR that is not a goal-directed task, relies primarily on hippocampal activity, and outcomes are highly dependent on the properties of the objects used [28].

Our results also underscore the potential of utilizing an appetitive-based maze under minimal food restriction to detect cognitive impairments at an early stage, preceding the onset of motor dysfunction. The longer time for completion and frequency of errors in PINK1^-/-^ rats indicate that this PD model may be deficient in spatial learning, egocentric navigation, and/or episodic memory in the premotor phase. The largest difference in scores between genotypes was on testing days 1 and 2. Considering the 2-day period between final training and start of the testing phase, the results suggest that 4-month-old PINK1^-/-^ rats have both learning and memory consolidation deficits – a neurological symptom that is not a characteristic observed in their age-matched WT counterparts. Although we identified overall differences in genotype performance, we did not identify clear learning acquisition to the maze path across the 5 consecutive trial days in either genotype. Considering the low number of errors made after the training phase, a more complex configuration of the maze may be able to discern differences in acquisition learning between the genotypes.

Cognitive decline in Parkinson’s disease (PD) represents an early symptom that significantly compromises the quality of human life, yet it remains largely unresponsive to currently available pharmacological treatments [2-4, 29]. A better understanding of the neuropathological changes that underlie such deficits will allow a path toward new treatment development and possible delay of disease progression. The major issue at present is to recognize that this specific decline occurs in the prodromal stage of the disease, manifesting in a heterogeneous manner as subtle alterations in executive functioning, working memory, attention, or visuospatial ability [2-3, 29-30]. Our implementation of the preclinical 5-T maze, under limited added stressors, such as an extended period of food restriction, appears to capture the underlying neurobiology of subtle decline in these specific cognitive domains. This increases the likelihood of capturing relevant neurobiological deficits in affected areas, such as the PFC, that are most responsive and most vulnerable to disease progression [19, 29-30]. Our results suggest that the PINK1^-/-^ rat can serve as an appropriate preclinical model for investigating the early neurobiological mechanisms associated with cognitive functioning also observed in human PD.

This is the first study, to our knowledge, to explore the PINK1^-/-^ rat’s capability to complete a food incentivized maze, during the premotor phase. In line with the findings of this study, prior cohorts of PINK1^-/-^ rats in our laboratory have consistently exhibited hyperkinetic behavior before the age of 6 months [8]. Notably, these outcomes contrast with reports from other studies, where the timing of initial motor deficits and the extent of dopamine (DA) loss in this model range widely, with some indicating no deficits and others observing deficits emerging between 4 to 8 months [7-9, 31]. This period of observed hyperkinesia may be attributed to a compensatory phase preceding symptomatic DA loss, potentially reflecting the premotor phase of Parkinson’s disease [32-33]. While other types of motor tests in rats may reveal specific deficits, such as in gait, the OFT we have used here assesses both spontaneous movement and speed, two motor aspects critical to maze completion. As PINK1^-/-^ rats exhibited a hyperkinetic phenotype, it is unlikely that motor dysfunction can explain their greater time taken to complete the 5-T maze.

While this study did not assess neurobiological differences associated with general, non-motor symptoms (NMS), brain imaging studies of PINK1^-/-^ rats have found decreased anisotropy in the olfactory system, hypothalamus, thalamus, nucleus accumbens, and cerebellum at postnatal weeks 12-13 [14]. A study using resting state functional MRI identified reduced connectivity between the neostriatum, midbrain DA regions, hypothalamus, and thalamus, and increased connectivity between ventral midbrain DA regions and hippocampus in 6-8 month-old PINK1-/-rats compared to WT [15]. Future studies are required to assess mechanisms in prefrontal cortex during this stage, as well as the progression of this cognitive phenotype, as some studies suggest that early cognitive decline may be indicative of disease trajectory [29]. Additionally, it would be beneficial to see if PINK1^-/-^ rats are able to navigate more complex paths in the maze which may better assess cognitive flexibility versus memory capacity. Finally, as this study served as a proof-of-concept study, only male rats were tested. Future studies will include evalution of sex differences to provide a more comprehensive assessment of this model. Overall, the results from this study show that male PINK1^-/-^ rats are compliant and capable of participating in a complex cognitive task yet show potential signs of prodromal cognitive decline as is seen in human PD.

## METHODS

PINK1^-/-^ (n=20) and wild-type (WT) Long Evans (n=14) rats were acquired from Envigo/Inotiv (Boyertown, PA) at 3-months-old and were handled daily for 1 week prior to any behavioral assessments. A power analysis established that this study possesses an 88% statistical power, given an alpha level of 0.05, to detect differences between WT and KO groups given the sample size. Rats were single-housed, kept on a 12-hr reverse light-dark cycle with *ad libitum* food and water. All procedures were approved by the Institutional Animal Care and Use Committee (IACUC) at the University of North Texas Health Science Center (UNTHSC) and the ACURO at the Department of Defense.

The CWM (MazeEngineers; Skokie, IL) is made of acrylic and consists of multiple arms arranged to produce a maze with different choice arms and one final endpoint. The maze was arranged into simplified “five choice arms” (5-T) following trial runs with pilot rats. The rats were tested on the maze one month after arrival to UNTHSC at 4 months of age. The task was conducted under regular light in the awake cycle in two phases: the training phase and the testing phase. During day one of the training phase, IACUC chow pellets were scattered inside the maze and rats were separately given 10 minutes to explore the area. On the second day, the quantity of scattered food was reduced and Stauffer’s animal crackers added to the endpoint. On the final day of training, all scattered food was removed, and crackers and chow pellets were placed exclusively at the final designated arm.

The 5-day testing phase was completed 2 days after the training phase. *Ad libitum* food from rats was reduced by 20% of regular intake starting after the training phase and maintained during the days of testing [14]. Rats were given 10 minutes to acclimate to the light and room condition before being placed in the maze. Rats were then individually video-recorded while the assessor remained in the room but away from the maze. Each rat was removed from the maze as soon as the camera screen showed it had reached the endpoint and collected food. Each rat was allowed a maximum of 5 minutes to complete the task and the maze was thoroughly cleaned with disinfectant wipes and dried between uses. Once returned to their cage, food trays were filled again to 80% of normal chow pellet consumption. Rats were scored on completion time and number of errors made. Errors constituted any time the rat entered an incorrect arm and/or on any incorrect turns made.

OFT chambers (Columbus Instruments Inc.; Columbus, OH) were used to measure spontaneous locomotor activity in the awake cycle. One hour of motor activity was quantified, both as average distance traveled and average speed, from three consecutive days one month pre-maze and one week post-maze (i.e. at 3 and 4 months-old) assessment to determine the impact of motor deficits on cognitive performance in completing the maze.

GraphPad Prism ver. 10 was used for statistical analyses. A student’s t-test was used to determine differences between genotypes on overall maze performance. A repeated measures two-way ANOVA was used to assess day to day performance in the maze considering the 5 consecutive days of testing. A two-way ANOVA with repeated measures was used to assess motor performance between PINK1^-/-^ and WT rats from arrival at 3 months to 4 months old. Grubb’s test and cut-off score criteria were used to identify any outliers.

## Funding

IS is supported by the National Institute on Aging Training Grant T32 AG 020494. The work was funded in part by the Department of Defense Parkinson’s Disease Research Program (Award W81XWH-19-1-0757) to MFS, an Institute for Healthy Aging at the University of North Texas Intramural Award to MFS and VAN, and an HSC Presidential Endowment to the Chair of Pharmacology & Neuroscience to DPS. We thank Dr. Jason R. Richardson for his support.

## Conflict of Interest

The authors have no conflict of interest to report.

## Data Availability Statement

The data supporting the findings of this study are available within the article.

## Notes

### Competing Interest Statement

The authors have declared no competing interest.

